# Correlation analysis of changes in the expression of *C1qtnf* superfamily genes in the hypothalamus, thymus, and lungs against the background of chronic social stress during the development of Lewis lung adenocarcinoma in mice

**DOI:** 10.64898/2026.07.02.735448

**Authors:** N.N. Kudryavtseva, D.A. Smagin, I.L. Kovalenko, N.A. Popova, M.B. Pavlova

## Abstract

It has been previously shown that chronic social defeat stress caused by paired agonistic interactions between male mice is accompanied by the development of depression-like state and immune deficiency. The aim of this study was to investigate changes in the expression of *C1qtnf* superfamily genes (encoding the complement component related with tumor necrosis factor) in the hypothalamus, thymus and lungs against the background of the Lewis lung adenocarcinoma growth. In the experiments, on the 5th day of social stress, male mice were injected with tumor cells into the tail vein. Chronic social stress continued for the next two weeks. The transcriptomes of the hypothalamus, thymus and lungs of mice were sequenced at the Genoanalytica Collective Center (http://genoanalytica.ru/, Moscow). Changes in the expression of the *C1qtnf* genes in the tissues of stressed mice were studied compared with the control and mice that were additionally injected with tumor cells. Overall, significant correlations were found between expression of most genes in each tissue of the experimental groups. In the hypothalamus of stressed animals, when tumor cells were introduced, an increase in the expression of the genes *C1qtnf1, C1qtnf2, C1qtnf3, C1qtnf6* and *C1qtnf7* was observed compared to controls. In the thymus of these animals, tumor cell injection increased expression of the *C1qtnf1, C1qtnf5*, and *C1qtnf6* genes. In the lung of tumor-injected stressed mice, expression of the *C1qtnf1, C1qtnf2, C1qtnf7*, and *C1qtnf9* genes was decreased relative to controls and non-tumor-injected depressed mice, reaching near-zero levels in some mice. Analysis of *C1qtnf* superfamily gene expression in the all tissues revealed negative correlations between the expression of the *C1qtnf1, C1qtnf2*, and *C1qtnf7* genes in the hypothalamus and lungs indicating synchronization of processes against the background of social stress and Levis lung adenocarcinoma.

## Introduction

Previous studies have shown that chronic social defeat stress caused by repeated negative social experiences in daily agonistic interactions leads to the formation of mixed anxiety/depression-like state in mice [Kudryavtseva et al., 1991; Kudryavtseva, 2021 review], which is accompanied by pronounced development of immune deficiency, manifested in a decrease in overall resistance, as well as changes in humoral and cellular immunity [Devoino et al., 1993; Popova et al., 1996; Gryazeva et al., 2001; Tenditnik et al., 2004; Kudryavtseva et al., 2011] and by increased oncogenic processes [Kudryavtseva et al., 2019, review], as evidenced by increased metastasis of Lewis lung adenocarcinoma in stressed animals compared to the control [Kaledin, Kudryavtseva, 1992; Kaledin et al., 2006; Kudryavtseva et al., 2007] (Fig. 1). These data were subsequently confirmed by changes in the expression of many carcinogenesis and apoptosis genes in the hypothalamus of these mice presented in [Galyamina et al., 2022].

**Fig. 1.**
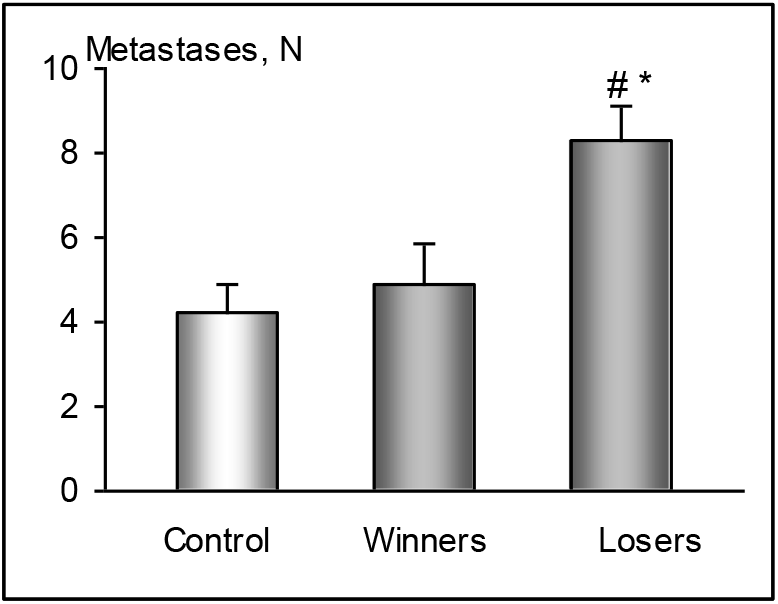
Lewis lung adenocarcinoma metastases (N) in male mice with positive (winners) and negative (losers) social experience in daily agonistic interactions. In this experiments tumor cells were injected into the tail vein of C57BL/6J mice [Kudryavtseva et al., 2007; Kaledin et al., 2009;]. * p < 0.01 - control *vs*. losers, stressed mice; - # p < 0.01-losers *vs*. winners, aggressive mice (t-Student test).

This study is focused on genes of the *C1qtnf* superfamily - *C1qtnf1, C1qtnf2, C1qtnf3, C1qtnf4, C1qtnf5, C1qtnf6, C1qtnf7, C1qtnf9*, which include the gene encoding the complement component (C1q) related with tumor necrosis factor (tnf). C1Q is a target protein of the classical complement pathway and a major link between innate and acquired immunity [Kishore et al., 2004].

Members of the C1Q and TNF family are thought to be involved in such diverse processes as inflammation, apoptosis, autoimmunity, cellular differentiation, organogenesis, and insulin-resistant obesity. The encoded C1QTNF proteins (CTRPs) are known to be members of the adiponectin family, which have attracted attention in recent years and to play an important role in the progression of many types of tumors, including liver, colon, and lung cancers through the activation of multiple signaling pathways [Li et al., 2017; Zhang et al., 2017; Kong et al., 2021; Lin et al., 2022; Zhang et al., 2023, reviews]. As a pattern recognition molecule of innate immunity, C1Q can interact with a wide range of ligands and modulate immune cells [Kishore et al., 2004].

In addition, complement activation is a component of innate immunity and a defense mechanism against invading pathogens. It is also involved in the adaptive immune response, inflammation, hemostasis, embryogenesis, repair and development of organs [reviewed in Afshar-Kharghan, 2017]. It is believed that the complement system of carcinogenesis can control tumor growth [Macor et al. 2018; Pio et al., 2014] and can be used for early diagnosis, prevention and development of innovative therapy, being its target [Yi et al., 2023; Merle, Roumenina, 2024]. It is argued that the interaction of complement components in each type of cancer is unique and is regulated by the properties of tumor cells. This concept is believed to be crucial for the development of effective therapeutic strategies [Roumenina et al., 2019]. At the molecular level, activation of complementary systems leads to changes in cellular signals, which can promote tumor cell proliferation. It is assumed that complementary interactions between cells and their environment can contribute to the formation of a microenvironment favorable for tumor growth. A decrease in the expression of complementary components can also lead to a weakening of the antimutagenic response, contributing to the progression of the tumor process.

In this work, we studied the change in the expression of the *C1qtnf* superfamily genes in the hypothalamus, thymus and lungs against the background of the administration of Lewis lung adenocarcinoma cells to mice under conditions of chronic social defeat stress caused by daily agonistic interactions with aggressive partners. In addition, the participation of the *Adipoq* gene was studied, in which the complementary component C1Q is associated with the collagen domain. This gene encodes the protein adiponectin, which is a hormone that has been shown to be involved in metabolic disorders [Tang et al., 2024; Zhang et al., 2023], which can lead, for example, to type 2 diabetes [Mamashli et al., 2025], obesity, and atherosclerosis [Chahirou et al., 2018]. People with lower adiponectin levels have reduced cognitive function [Feinkohl et al., 2024]. Its participation in the mechanisms of proliferation, apoptosis, and autophagy of glioma cells has also been shown [Sun et al., 2024] as well as in modulating the development of osteosarcoma [Zhang et al., 2025] and hepatocellular carcinoma [Yao et al. 2023; Yu et al., 2024].

It has also been shown that the state of the immune system is of great importance for tumor development. Understanding the mechanisms of *C1qtnf* genes involvement in tumor formation is believed to help in the development of new cancer treatments, and the proteins they encode could be considered as diagnostic markers and therapeutic targets for numerous types of cancer [Kong et al., 2021; Lin et al., 2022; Zhang et al., 2023].

The aim of this work was to study the changes in the expression of *C1qtnf* superfamily genes in the hypothalamus and peripheral tissues - thymus and lungs - both under the influence of chronic social stress, accompanying by development of a depression-like state in mice, and against the background of the administration of Lewis lung adenocarcinoma cells into stressed animals during the formation of depressive pathology. We believe that the experimental approach we used to study the changes of expression of *C1qtnf* superfamily genes against the background of the development of psychogenic immunodeficiency, tumor processes can open up prospects for the development of new strategies for the treatment of tumors. The task also included the identification of correlative relationships between changes in gene expression in every tissue (hypothalamus, thymus, lungs) and those common to different tissues.

## Materials and methods

### Animals

The experiments were performed on male C57BL/6J mice aged 2.5 months and weighing 26–28 g. The animals were brought from the Laboratory Animal Nursery of the Institute of Bioorganic Chemistry of the Russian Academy of Sciences (Pushchino, Moscow Region). The animals had constant unlimited access to food (pellets) and water and were kept in a 12-hour light/dark cycle. All procedures were carried out in accordance with international guidelines for animal experiments (European Communities Directive (86/609 EEC).

### Chronic social conflict model

Chronic social stress in mice was formed using the chronic social conflict model [Kudryavtseva et al., 2014; Kudryavtseva, 2021]. In daily agonistic interactions, animals acquired chronic negative experience of social defeats with an aggressive partner (winners) and living with him in conditions of constant sensory contact. For this, animals were placed in pairs in experimental cages divided in half by a transparent partition with holes, which allowed the mice to see, hear, and smell each other (sensory contact), but prevented physical interaction. Every day in the afternoon (14:00–17:00), the partition was removed, which led to agonistic interactions. The interaction of males was stopped if intense attacks from the attacking individuals lasted more than three minutes (sometimes less), installing a partition between them to avoid physical damage. During the first 2-3 days, the tests revealed winners who demonstrated aggression towards the partner, and males who suffered defeats when interacting with the aggressive partner. Subsequently, every day after the test, the defeated male was transferred to other cage with an unfamiliar aggressive partner located behind the partition.

For neurogenomic studies, animals with a pronounced phenotype of depression-like behavior (hereinafter, losers, depressed mice) formed in daily agonistic interactions, and demonstrating a decrease in motor activity, avoidance of social contacts, and increased anxiety in various behavioral tests were used. The following groups were studied in the experiment: 1) control animals - individuals without repeated experience of agonistic interactions; 2) depressed mice after 20 days of agonistic interactions; 3) depressed males, which on the fifth day of agonistic interactions the Lewis lung adenocarcinoma cells (1 million tumor cells in 0.1 ml of physiological solution) were transplanted into the tail vein (hereinafter referred to as depressed tumor-injected mice). Agonistic interactions were then continued for another 14 days. The behavior of the animals was registered daily in the experimental protocols.

Animals of the experimental groups were studied the day after the last confrontation, simultaneously with control mice. The hypothalamus, thymus and lungs were isolated from all mice. The hypothalamus was removed in accordance with the anatomical atlas of the brain (Allen Mouse Brain Atlas; http://mouse.brain-map.org/static/atlas). All samples were placed in RNAlater solution (Life Technologies, USA) and stored at - 70 °C until sequencing.

### RNA-Seq analysis

Samples of the hypothalamus, thymus and lungs were sequenced at the Genoanalytica Collective Center (http://genoanalytica.ru, Moscow, Russia). The quality and quantity of the isolated total RNA were assessed using a BioAnalyzer with the RNA 6000 Nano Kit (Agilent). The poly(A) RNA fraction was then purified using oligo(dT) magnetic beads (Dynabeads® mRNA Purification Kit, Ambion) according to the manufacturer’s instructions. cDNA libraries were constructed using the NEBNext protocol for Illumina (NEB, USA). Sequencing of the cDNA libraries was performed on the Illumina Hiseq 1500 platform (Illumina Sequencing, USA). Six animals were analyzed in each group. All samples were analyzed separately. The sequencing depth was more than 20 million reads per sample. Genes were considered differentially expressed if their expression levels differed between mice injected with and without tumor cells compared to control animals (p < 0.05). The following genes were taken into account: C1qtnf1, C1qtnf2, C1qtnf3, C1qtnf4, C1qtnf5, C1qtnf6, C1qtnf7, C1qtnf9, and Adipoq. Genes with low (< 10.00 - normalized read count), hereinafter RPKM (Reads Per Kilobase per Million Mapped Reads), as well as genes that did not differ in expression levels when comparing mice from all experimental groups were not taken into account in animals. The decision on statistically significant differences in expression was also made using the correction for multiple comparisons (q-values are adjusted p-values according to the Benjamini-Hochberg (FDR). The results were visualized using the SRplot bioinformatics resource (https://www.bioinformatics.com.cn/en) [Tang et al., 2023] and XLSTAT Version 2016.02.28451 (https://www.xlstat.com). Additionally, correlation analysis of gene expression in animals in different tissues was also used.

Functional annotation of differentially expressed genes (DEGs) was performed using the DAVID Bioinformatics Resources 6.7 Internet resource (http://david.abcc.ncifcrf.gov), as well as the KEGG pathway (https://www.genome.jp/kegg/pathway.html) and Gene Cards databases (https://www.genecards.org).

## Results

Expression of genes was compared across the experimental groups selected for analysis: 1) control mice *vs*. depressed mice; 2) control mice *vs*. depressed mice with the injection of tumor cells (hereinafter, depressed tumor-injected mice); 3) depressed mice *vs*. depressed tumor-injected mice. Means and SEM are presented for 6 mice in every group and tissue in Table 1 and Supplementary Table 1. Figures 2, 3, 4 demonstrate visualization of the expression for DEGs in the hypothalamus, thymus and lung in mice.

**Table 1.** The *C1qthf* superfamily DEGs when comparing the experimental groups.

| Table 1. The *Clqthf* superfamily DEGs when comparing the experimental groups |  |  |  |
| --- | --- | --- | --- |
| DEGs | C vs D | C vs DT | D vs DT |
| **Hypothalamus** |  |  |  |
| *Clqtnf1* | — | ▲▲** | ▲▲▲** |
| *Clqtnf2* | — | ▲▲▲** | ▲▲▲** |
| *Clqtnf3* | — | ▲ | — |
| *Clqtnf6* | — | ▲▲* | ▲▲** |
| *Clqtnf7* | — | ▲▲▲*** | ▲▲▲*** |
| *Adipoq* | — | ▲▲▲*** | ▲▲* |
| **Thymus** |  |  |  |
| *Clqtnf1* | — | ▲▲ | ▲▲* |
| *Clqtnf5* | — | ▲ | ▲▲▲** |
| *Clqtnf6* | — | ▲▲▲*** | ▲▲▲*** |
| *Adipoq* | ▲ | — | ▲ |
| **Lung** |  |  |  |
| *Clqtnf1* | — | ▼ | ▼ |
| *Clqtnf2* | — | ▼ | ▼ |
| *Clqtnf4* | — | ▲▲▲*** | ▲▲▲* |
| *Clqtnf7* | — | ▼ | — |
| *Clqtnf9* | — | ▼ | ▼ |
| *Adipoq* | — | — | ▼ |
| ▲/▼ p < 0.05; ▲▲ p < 0.01; ▲▲▲ p < 0.001 |  |  |  |
| * - q < 0.05; ** - q < 0.01; *** - q < 0.001 |  |  |  |
| C – control; D – depressed mice; DT – depressed tumor-injected mice. Significant differences are presented for 6 mice in every group and tissue in Supplementary Table 1 |  |  |  |

**Fig. 2.**
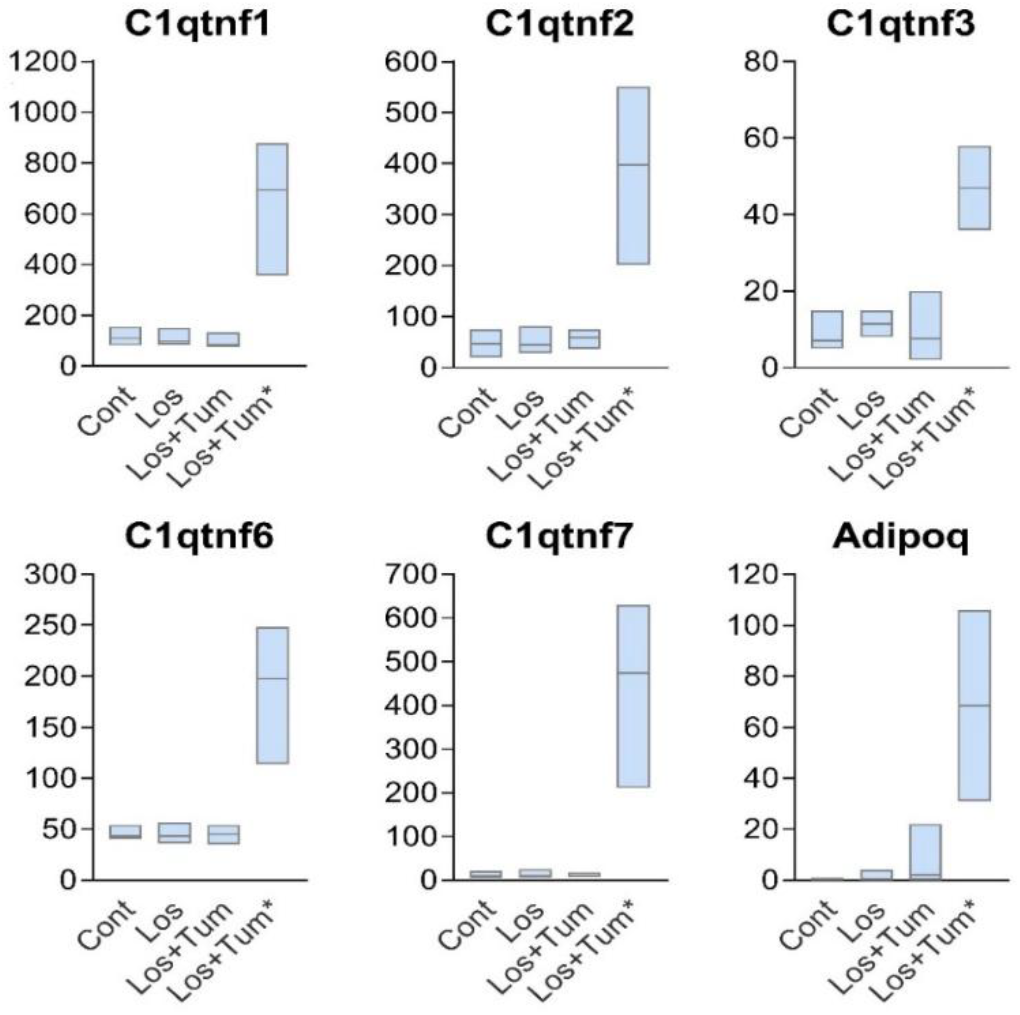
Visualization of expression for DEGs in the hypothalamus of different experimental groups in mice: Cont – control, intact mice; Los – losers, depressed mice; Los+Tum – losers, depressed tumor-injected mice; Los+Tum* – losers, depressed tumor-injected mice with maximal changes in gene expression. Data is presented as a floating bar (min to max value) with a line indicating the median level. Significant differences between groups of comparison are presented in Table 1.

**Fig. 3.**
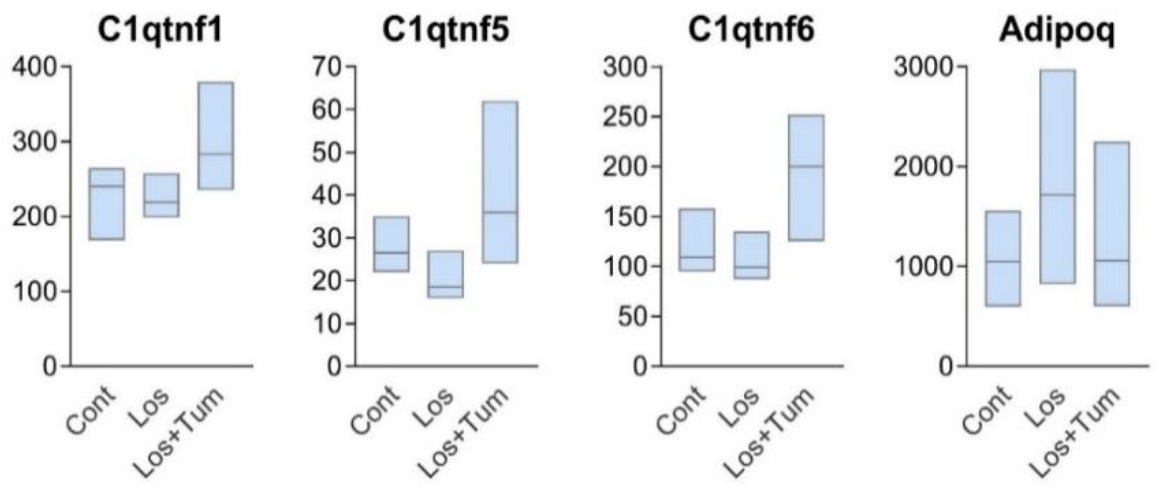
Visualization of expression for DEGs in the thymus of different experimental groups in mice: Cont – control, Los – losers, depressed mice; Los+Tum – losers, depressed tumor-injected mice. Data is shown as a floating bar (min to max value) with a line indicating the median level. Significant differences between groups are presented in Table 1 and Supplementary Table 1.

**Fig. 4.**
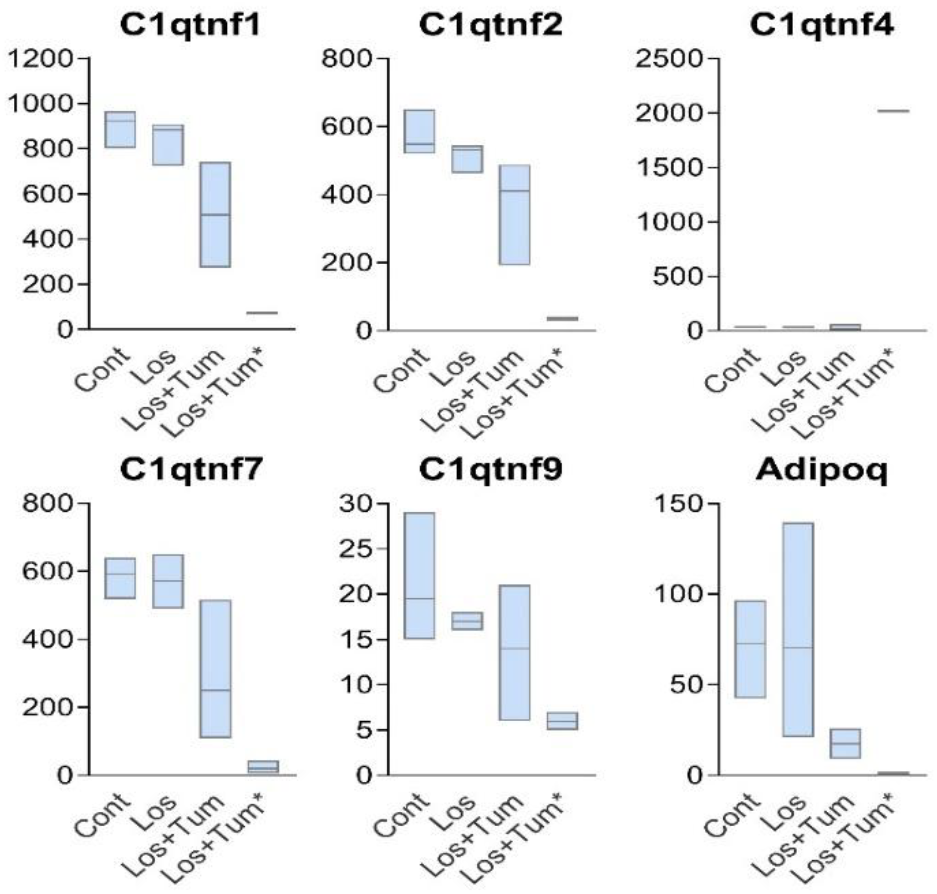
Visualization of expression for DEGs in the lung of different experimental groups in mice: Cont – control, Los – losers, depressed mice; Los+Tum – losers, depressed tumor-injected mice; Los+Tum* – losers, depressed mice with maximal changes in group depressed tumor-injected mice. Data is presented as a floating bar (min to max value) with a line indicating the median level. Significant differences between groups are presented in Table 1 and Supplementary Table 1.

It should be noted that the same gene could have different levels of expression in tissues, and at the same time, different genes could differ in the direction of altered expression in different tissues. As example (Supplementary Table 1): expression of *C1qtnf1* gene in the controls was 115.30 ± 11.01 RPKM in the hypothalamus; 234.50 ± 14.69 RPKM – in the thymus and 902,30 ± 24.15 RPKM in the lung. Expression of the *Adipoq* gene in the controls was about 0.0 – in the hypothalamus, 1038.00 ± 150.30 – in the thymus and 71,80 ± 9.37 – in the lung.

By analyzing data from the hypothalamus and lungs, but not the thymus, we identified experimental mice that differed significantly in the extent to which tumor-induced changes in the expression of certain genes occurred. This led to the need to isolate among the animals with the injection of tumor cells two separate groups: Los+Tum and Los+Tum* groups, in which the greatest differences in the severity and direction of changes in gene expression were found (Fig. 2 and Fig. 4).

### Changes in the expression of C1qtnf superfamily genes in depressed mice under the influence of tumor growth in the hypothalamus

In the hypothalamus, the initial values of gene expression in the control animals (C) were relatively low, with the exception of *C1qtnf5* (> 350 RPKM). However, expression of this gene was similar in all groups. The *Adipoq* gene had zero or very low expression values (Fig. 2, Supplementary Table 1, 2), which indicated that this gene in the hypothalamus of intact animals can most likely be attributed to the group of predominantly with low expression genes, like the *C1qtnf3* and *C1qtnf7*genes. Expression of these genes were increased only under tumor growth (DT group; Table 1, Supplementary Table 1). Visualization of the expression for each gene in the hypothalamus of mice of different experimental groups is presented in Fig. 2.

**Table 2.** Correlation relationships of expression of the *C1qtnf* superfamily genes in the hypothalamus of all groups of comparison.

|  | *Clqtnf1* | *Clqtnf2* | *Clqtnf3* | *Clqtnf6* | *Clqtnf7* | *Adipoq* |
| --- | --- | --- | --- | --- | --- | --- |
| *Clqtnf1* |  | 0.989*** | 0.928*** | 0.989*** | 0.996*** | 0.847*** |
| *Clqtnf2* | 0.989*** |  | 0.880*** | 0.979*** | 0.991*** | 0.785*** |
| *Clqtnf3* | 0.928*** | 0.880*** |  | 0.932*** | 0.915*** | 0.943*** |
| *Clqtnf6* | 0.989*** | 0.979*** | 0.932*** |  | 0.992*** | 0.881*** |
| *Clqtnf7* | 0.996*** | 0.991*** | 0.915*** | 0.992*** |  | 0.840*** |
| *Adipoq* | 0.847*** | 0.785*** | 0.943*** | 0.881*** | 0.840*** |  |
| n = 18; *** - p < 0.001 |  |  |  |  |  |  |

When comparing the ***Control mice vs. Depressed mice***, no genes were identified whose expression was significantly altered in the hypothalamus of depressed mice (C *vs* D, Table 1, Supplementary Table 1). The *C1qtnf3* and *Adipoq* genes had low values (< 10.00 RPKM on average, Supplementary Table 1, Fig. 2). When comparing the ***Control mice vs. Depressed tumor-injected mice*** (C *vs* DT), the expression of the *C1qtnf1, C1qtnf2, C1qtnf3, C1qtnf6, C1qtnf 7* and *Adipoq* genes was higher compared to the control (Table 1; (Los+Tum)+(Los+Tum*)). In two or three depressed tumor-treated mice, the gene expression was significantly higher, which allowed us to classify them into the Los+Tum* group (Fig. 2). When comparing the ***Depressed mice vs. Depressed tumor-injected mice***, it was shown that all genes - *C1qtnf1, C1qtnf2, C1qtnf6, C1qtnf7* and *Adipoq* - in depressed ***tumor-injected*** mice (Los+Tum)+(Los+Tum*) (D vs DT) had increased expression when compared to the group of depressed mice without tumor cells injection.

Correlation analysis of three comparison groups of mice (control, depressed without and with the tumor-injected cells) in the hypothalamus revealed a relationship between the expression of almost all genes with a high level of significance (Table 2). Moreover, all genes had a positive correlative relationship in the hypothalamus.

### Changes in the expression of C1qtnf superfamily genes in the thymus of depressed mice under the influence of tumor growth

In the thymus of control animals, the highest expression was observed for *C1qtnf1* (> 230 RPKM) and *Adipoq* (> 1000 RPKM) genes (Fig. 3, Supplementary Table 1). Visualization of expression for each gene in the mice from different experimental groups is presented in Fig. 3.

When comparing the ***Control mice vs. Depressed mice***, no genes were identified whose expression was significantly altered in the thymus of depressed mice (C *vs* D), except *Adipoq* gene: expression was higher in depressed mice (Table 1). When comparing the ***Control mice vs. Depressed tumor-injected mice***, expression was increased for the *C1qtnf1, C1qtnf5*, and *C1qtnf6* genes relative to the control. When comparing the ***Depressed mice vs. Depressed tumor-injected mice***, a significant increase in the expression of the *C1qtnf1, C1qtnf5*, and *C1qtnf6* genes was found (Table 1).

Correlation analysis of gene expression in the groups (control, depressed without and with the tumor cells injection) of comparison in the thymus revealed a relationship between changes in the expression of the C*1qtnf1, C1qtnf5* and *C1qtnf6* genes (but not the *Adipoq* gene), with a high degree of significance (Table 3).

**Table 3.** Correlation relationships of the expression of the *C1tnf* superfamily genes in the thymus of all groups of comparison.

| Thymus |  |  |  |  |
| --- | --- | --- | --- | --- |
|  | *Clqtnf1* | *Clqtnf5* | *Clqtnf6* | *Adipoq* |
| *Clqtnf1* |  | 0.637** | 0.761*** | 0.229 |
| *Clqtnf5* | 0.637** |  | 0.863*** | 0.096 |
| *Clqtnf6* | 0.761*** | 0.863*** |  | -0.020 |
| *Adipoq* | 0.229 | 0.096 | -0.020 |  |
| n = 18; ** p < 0.01; *** p < 0.001 |  |  |  |  |

### Changes in the expression of C1qthf superfamily genes in the lung of depressed mice under the influence of tumor growth

In the lung, in mice of the control group, the gene expression was initially significantly higher compared to other tissues for *C1qtnf1* (> 900 RPKM), *C1qtnf2* (> 500 RPKM) and *C1qtnf7* (about 600 RPKM) genes. At the same time, a large variability of data on the change in gene expression in the lung of depressed mice was found. As a result, similar with the hypothalamus, 2 groups of mice were isolated: The Los+Tum group and Los+Tum* group (Fig. 4). Large differences in the decrease in gene expression in the Los+Tum* group of mice compared to the Los+Tum group were observed for *C1qtnf1, C1qtnf2, C1qtnf7, C1qtnf9* and *Adipoq* genes.

When comparing the ***Control mice vs. Depressed mice*** (Los+Tum)+(Los-Tum*) (Table 1, Supplementary Table 1), there were no differences in expression levels. When comparing the ***Control mice vs. Depressed tumor-injected mice*** in both groups in total, a decrease in expression of the *C1qtnf1, C1qtnf2, C1qtnf7*, and *C1qtnf9* genes was observed. The pronounced decrease of expression of *Adipoq* gene, close to zero values was showed (Fig. 4). When comparing the ***Depressed mice vs. Depressed tumor-injected mice***, a decrease in expression of the *C1qtnf1, C1qtnf2, C1qtnf7, C1qtnf9*, and *Adipo*q genes was revealed compared to mice in the (Los+Tum)+(Los+Tum*) groups. Significantly increased expression was noted – about 2000 RPKM -for the *C1qtnf4* gene in two Los+Tum* mice (Fig. 4, Table 1).

Correlation analysis of gene expression in the groups (control, depressed without and with the tumor-injected cells) of comparison in the lung revealed a relationship between changes in the expression of the *C1qtnf1, C1qtnf2, C1qtnf7* genes with a high significance (Table 4). Significant correlations of genes with the expression level of the *Adipoq* gene and *C1qtnf9* were found (Table 4). Expression of the *C1qtnf4* genes negatively correlates with expression of the *C1qtnf1, C1qtnf2*, and *C1qtnf7* genes.

**Table 4.** Correlation relationships of the expression of *C1tnf* superfamily genes in the lung of all groups of comparison.

| Lung |  |  |  |  |  |  |
| --- | --- | --- | --- | --- | --- | --- |
|  | *Clqtnf1* | *Clqtnf2* | *Clqtnf4* | *Clqtnf7* | *Clqtnf9* | *Adipoq* |
| *Clqtnf1* |  | 0.944*** | -0.749*** | 0.613** | 0.466 | 0.234 |
| *Clqtnf2* | 0.944*** |  | -0.611** | 0.540* | 0.400 | 0.169 |
| *Clqtnf4* | -0.749*** | -0.611** |  | -0.604** | -0.320 | -0.171 |
| *Clqtnf7* | 0.613** | 0.540* | -0.604** |  | 0.396 | 0.196 |
| *Clqtnf9* | 0.466 | 0.400 | -0.320 | 0.396 |  | 0.927*** |
| *Adipoq* | 0.234 | 0.169 | -0.171 | 0.196 | 0.927*** |  |
| n = 18; * p < 0.05; ** p < 0.01; *** p < 0.001 |  |  |  |  |  |  |

### Principal component analysis for the lung

Principal components analysis (Fig. 4) of gene expression in the three groups (C - control, D - depressed mice and DT - depressed tumor-injected mice revealed that two of them - C (green) and D (blue) - have similar expression profiles excluding D4, while the DT (red) is significantly different and characterized by greater heterogeneity. The main differences between the groups are explained by the first principal component, indicating the presence of unique gene expression patterns in DT. The density of clusters C and D suggests that gene expression in these two groups is relatively stable and similar. DT mice with tumor injection stands out: it is likely that unique genes are activated or suppressed in the lung.

**Fig. 4.**
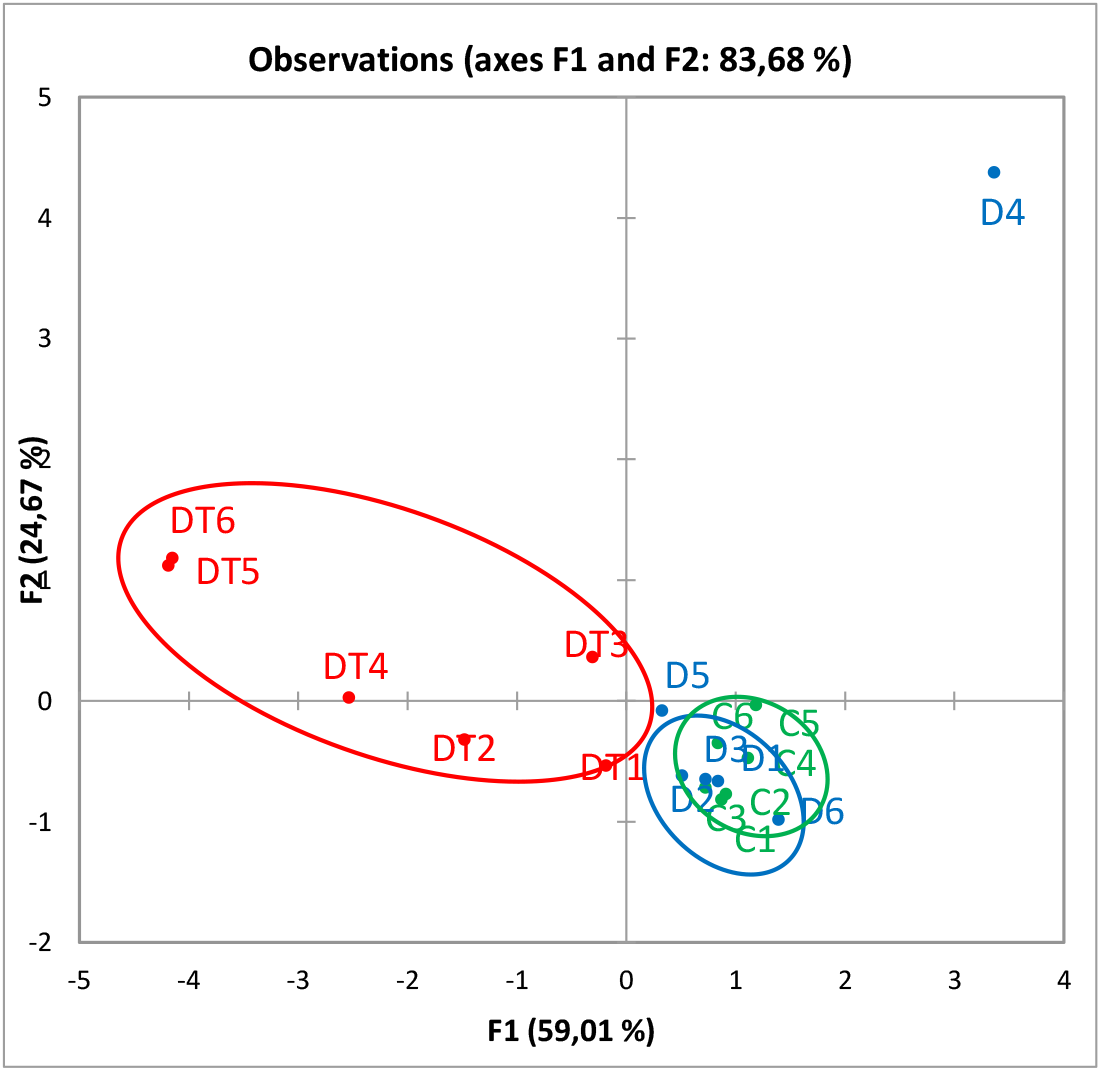
Principal components analysis (PCA) for 18 samples of DEGs in the lung of mice of different experimental groups: green C (1-6) – control, intact mice; blue D (1-6) – depressed mice; red DT (1-6) – depressed tumor-injected mice.

#### Correlation analysis of the same genes expression in different tissues

Changes of expression of the *C1qtnf1* and *Adipoq* genes were revealed in all tissues (Table 5). In the hypothalamus and lung – the *C1qtnf2* and *C1qtnf7* genes, and in the hypothalamus and thymus – the *C1qtnf6* gene were changed its expression. The *C1qtnf3* expression was changed only in the hypothalamus, and the *C1qtnf4* and *C1qtnf9* genes – only in the lung, and *C1qtnf5* gene – only in the thymus.

**Table 5.**
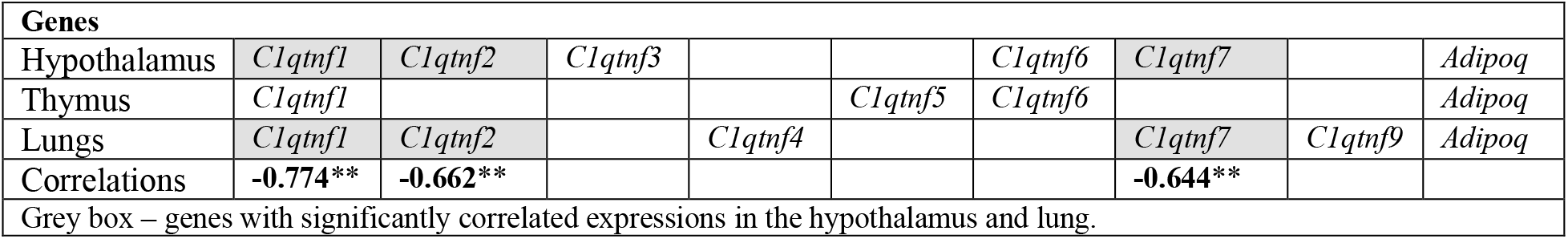
The *C1qtnf* superfamily genes changing the expression in different tissues under the influence of experimental exposure.

| Genes |  |  |  |  |  |  |  |  |  |
| --- | --- | --- | --- | --- | --- | --- | --- | --- | --- |
| Hypothalamus | *Clqtnf1* | *Clqtnf2* | *Clqtnf3* |  |  | *Clqtnf6* | *Clqtnf7* |  | *Adipoq* |
| Thymus | *Clqtnf1* |  |  |  | *Clqtnf5* | *Clqtnf6* |  |  | *Adipoq* |
| Lungs | *Clqtnf1* | *Clqtnf2* |  | *Clqtnf4* |  |  | *Clqtnf7* | *Clqtnf9* | *Adipoq* |
| Correlations | -0.774** | -0.662** |  |  |  |  | -0.644** |  |  |

Grey box – genes with significantly correlated expressions in the hypothalamus and lung.

Significant negative correlations of changes in the expression of the same genes in different tissues were found in the hypothalamus and lungs for the *C1qtnf1* genes (-0.774), *C1qtnf2* (-0.662), and *C1qtnf7* (-0.644). There were no significant correlations of the changed expression of other genes - between the hypothalamus and thymus, as well as between the thymus and lung.

## Discussion

The complementary components (C1q) attracted attention in connection with the discovery of their great role in the development of cancer and not only [Afshar-Kharghan et al., 2017; Ajona et al., 2019; Qiu et al., 2023; Zhan et al., 2023]. It is believed that at the molecular level, activation of complementary systems leads to a change in cell signals that can contribute to the proliferation of tumor cells [Roumenina et al., 2019; Merle, Roumenina, 2024, reviews]. At the same time, a decrease in the expression of genes encoding tumor necrosis factors (TNF) is an important area of biomedical research aimed at understanding the mechanism of development of various diseases, including cancer and inflammatory disease. These factors also play a key role in the immune processes, regulation of cellular growth and differentiation, as well as in the induction of apoptosis of malignant cells [Afshar-Kharghan, 2017; Ajona et al., 2019; Berraondo et al., 2016; Kemper, Köhl, 2018]. However, all studies note that understanding of developing processes of cancer depends on many components.

This study draws attention to the genes of *C1qtnf* superfamily encoding the complementary components of C1q, related with a tumor necrosis factor (tnf) in the hypothalamus, thymus and lungs of mice with a depression-like state formed under the influence of chronic social stress during of the growth of Lewis lung adenocarcinoma. The response to chronic social stress and/or tumor cell injection has been shown to vary depending on the gene and tissue (hypothalamus, thymus, lung). A gene may not respond to stimuli in one tissue, but change its expression in another, and the degree of expression of the changes may vary: we believe this is due to the different influence of the biochemical and cellular environment, and may also depend on the inclusion of its proteins in different neural networks.

Terms assessing the functional state of genes have been used previously, including in the context of cancer development [Meyerson, 2007; Rich et al., 2024; Zhang X. et al., 2025]. We found it possible to apply some of these terms to interpret changes in the state of *C1qtnf* genes under the influence of experimental interactions (chronic social stress and tumor cell injection). Analysis of the results allowed initially to identify genes that respond differently to the experimental exposure and to suggest their different states.

**In the hypothalamus**, expression of all *C1qtnf* genes was similar in the control and depressed mice. These genes were assigned to the group of “non-working” or “silent” genes in this tissue. However, its expression in the depressed tumor-injected mice [(Los+Tum)+(Los+Tum*)] was significantly higher (Fig. 2, Table 1, Supplementary Table 1), especially in the Los+Tum* group. Since neurophysiological events under the influence of chronic stress occur initially in the hypothalamus, then it is reasonable to assume that proteins encoded by the *C1qtnf* genes in this brain region may serve as markers of susceptibility to increased carcinogenesis.

**In the thymus**, the *C1qtnf1, C1qtnf5*, and *C1qtnf6* genes significantly increase the expression of genes in depressed tumor-injected mice compared to depressed mice and control (Fig. 3, Table 1, Supplementary Table 1) and may be classified as “Cancer genes**”** involved in the development of the tumor process. If taking into account the activation of the expression of these genes in response to a decreased immunity, as was previously shown in depressed mice under the influence of chronic social stress [Devoino et al., 1993; Popova et al., 1996; Tenditnik et al. 2004; Gryazeva et al., 2001; Kudryavtseva et al., 2011; Kudryavtseva et al., 2019], it can be assumed that in the thymus, the proteins encoded by these genes may serve as peripheral markers of predisposition to increased carcinogenesis as was shown earlier [Kaledin, Kudryavtseva, 1992; Kaledin et al., 2006; Kudryavtseva et al., 2007]. It should be noted that the increase in gene expression, at least, for *C1qtnf1* and *C1qtn6* genes in the thymus under the influence of the tumor growth is similar to that in the hypothalamus.

**In the lungs**, a different picture is observed. Gene expression in the Los+Tum group (*C1qtnf1, C1qtnf2, C1qtnf9*) is reduced and can be considered as “broken” (damaged). Meanwhile, in mice from the Los+Tum* group, the expression of these genes is virtually zero (Fig. 4, Table 1, Supplementary Table 1). These genes can be tentatively classified as “dead” or “dying” genes influenced by tumor growth. However, it should be noted that these terms require confirmation at the genomic and molecular levels.

The exception is the *C1qtnf4* gene, which is a unique member of the *C1qtnf* superfamily genes. In mice of the Los+Tum* group this gene is activated, reaching the highest values of expression compared even with genes in other tissues - thymus and hypothalamus. The protein encoded by the *C1qtnf4* gene is considered as a potential cytokine that promotes the survival of human cancer cells [Li et al., 2011]. According to some data [Vester et al., 2021], the *C1qtnf4* gene encodes a protein that has structural homology with both of *C1q* and the superfamily of tumor necrosis factors. For this gene, significant negative correlations with the *C1qtnf1, C1qtnf2*, and *C1qtnf7* genes are installed (Table 4) in the lung.

The correlation analysis of changed genes expression in all groups of comparison (control, depressed mice and depressed tumor-injected mice) revealed the relationship of changes in three tissues with high significances. In the hypothalamus, positive correlations were found between all genes. In the thymus, significant correlation relationships were installed for *C1qtnf1, C1qtnf5* and *C1qtnf6* genes. In the lung, the correlation analysis of three groups of comparison revealed a positive relationship of changes in the expression of *C1qtnf1* and *C1qtnf2* and *C1qtnf7*, as well as *Adipoq* and *C1qtnf9* genes. Only for *C1qtnf4* gene, a negative relationship with the expression of the *C1qtnf1, C1qtnf2* and *C1qtnf7* genes was established. The unidirectional changes in the expression of genes indicate the influence of a unified exposure and uniformity of changes in the expression of genes involved in the development of depressed mice and depressed tumor-injected mice. A study of correlations between changes in the expression of the same genes in the hypothalamus and peripheral tissues revealed significant negative correlations in the hypothalamus and lungs for the *C1qtnf1, C1qtnf2*, and *C1qtnf7* genes. This suggests that increased gene expression in the hypothalamus is associated with decreased expression in the lung.

Particular attention attracts the *C1qtnf1* gene, which demonstrates changes in all three tissues, and allows to consider it as a key gene that plays the main role in the development of cancer processes. It is this gene, according to available data, that is involved in the development of various types of cancer, in particular kidney cancer [Qiu et al., 2023], hepatocellular carcinoma [Lin et al 2021; Yu et al., 2024]; malignant colorectal cancer [Jin et al., 2020]; osteosarcoma [Zhang et al., 2025] and others.

The significant correlations between the same genes in different tissues confirms the previously advanced our hypothesis described in the article [Galyamina et al., 2022] which suggests that the detected changes in the expression of carcinogenesis and apoptosis genes in the hypothalamus of depressed mice can reflect changes in peripheral tissues, in particular, in tumors, and *vice versa*. Such mechanism is possible if we assume that there is coordination of these processes in the brain and peripheral tissues (for example, in the lung) and if changes in genes expression induce the same factors. For example, it can be corticosteroids, other hormones or neurotransmitters in the blood, caused, in our case, chronic social stress.

This preliminary conclusion was confirmed previously. A comparative analysis of altered gene expression in the hypothalamus of depressed animals and key genes in lung cancer patients revealed similarities in several genes, particularly the Eno2 gene [Galyamina et al., 2024]. These findings were indirect evidence that in the brain tissue, for example, in the hypothalamus, altered expression of genes can to a certain extent correspond to changes in peripheral tissues. Our study confirmed this hypothesis by identifying simultaneous changes in the expression of genes directly linked to carcinogenesis in the hypothalamus, thymus, and lungs. The majority of significant correlations between the same genes, *C1qtnf1, C1qtnf2*, and *C1qtnf7*, were found in the hypothalamus and lungs.

It is also necessary to take into account the results indicating that the psychoemotional status through the modification of immune reactivity affect the growth and metastasis of the tumor in mice [Kudryavtseva et al., 2019, review]. Therefore, a possible explanation of changes in the lungs of depressed mice in our experiments may be a decreased immunity as the main factor that determines changes in expression of the genes in these experiments. Some neuroendocrine changes can act as peculiar growth factors and modify the development of tumors and metastasis both by direct action on the tumor cells, and indirectly through the effect on the vascular direction of target organs. Physiological features such as the activity of tissue macrophages and NK cells, endothelial capillary permeability, vascularization rate, etc. can influence the colonization of target organs by tumor cells can affect colonization with tumor cells in target organs [Spiegel, Sephton, 2001; Rowse et al., 1992; Palermo-Neto et al., 2003].

In addition, there were a number of results indicating that the enhance of tumor metastasis is stopped by antagonists of the receptors of some neurotransmitters [Giraldi et al., 1994; Perissin et al., 1996; Beneliyahu et al. 2000; Freire-Garabal et al., 2004; Palm et al., 2006]. Our results on the pharmacological correction of emotional status using anxiolytic diazepam [Kaledin et al. 2009; Tenditnik et al., 2010] or ethanol [Ilnitskaya et al., 2009], which reduced the level of anxiety in depressed male mice, preventing the development of immune deficiency, indicated a close relationship of these processes. At the same time, the activation of the immune system using the chronic administration of roncoleukin did not affect this process [Shurlygina et al., 2015].

Recent data [Juszczak et al., 2025] confirmed our results and the hypothesis about the possible correlation of the expression of the same genes in the brain and peripheral tissues. These authors conducted a large experiment to check the effects of long-term exposure to corticosterone on the hippocampus transcript (RNA sequencing). Despite the difficulties of interpretation, as the authors write, the results indicated that prolonged consumption of corticosterone, which to some extent imitated the effects of chronic stress, leads to a large set of transcription effects in the hippocampus [Juszczak et al., 2025].

Changes in gene expression in mice during the development of a depressive-like state and decreased immunity, accompanied by changes in the expression of apoptosis and carcinogenesis genes in the hypothalamus under the influence of chronic social stress, suggest that corticosteroids may be the main stimulators of correlative changes in gene expression in the brain and peripheral tissues. Further studies in this area are necessary for the development of new therapeutic approaches, which are aimed at specific manipulation of genes expression encoding tumor necrosis factors, that can increase the effectiveness of treatment and minimize side effects in patients.

## Limitations

In this study, attention was paid to changes in the expression of the *C1qtnf* superfamily genes, which work in ensemble of the complementary component C1q with tumor necrosis factor (Tnf). However, it is also necessary to take into account that in addition to these genes, there is a superfamily of 19 genes encoding tumor necrosis factor (*Tnf*), as well as a superfamily of 29 *TnfR* genes encoding receptors [https://en.wikipedia.org/wiki/tnf_receptor_superfamily [Locksley et al., 2001; Hehlgans, Pfeffer, 2005; Zhang et al., 2021]. These genes can undoubtedly be involved in the process of carcinogenesis, and are related with immune, inflammatory, and cellular processes.

## Supporting information

Supplementary Table S1. The C1qtnf DEGs in the hypothalamus, thymus and lung

## Supplementary Materials

Supplementary Table 1: RPKM values for C1qtnf DEGs in the hypothalamus, lungs, thymus.

## Author Contributions

Conceptualization, N.N.K.; data curation, I.L.K., D.A.S.; formal analysis, N.N.K., D.A.S., M.B.P.; investigation, D.A.S., N.A.P., M.B.P., N.N.K.; methodology and supervision, N.N.K.; writing—original draft, N.N.K., D.A.S.; writing—review and editing N.N.K., D.A.S., M.B.P.

All authors have read and agreed to the published version of the manuscript.

The study is supported by State Funding of the I.P. Pavlov Institute of Physiology, Russian Academy of Sciences

(*N*o1021062411629-7-3.1.4).

## Institutional Review Board Statement

All procedures were conducted in compliance with European Communities Council Directive 210/63/EU of 22 September 2010. The study protocol was approved by Scientific Council No. 9 of the Institute of Cytology and Genetics SB RAS on 24 March 2010, No. 613 (Novosibirsk, Russia).

## Data Availability Statement

All relevant data are available in the Supplementary Materials and from the authors.

## Acknowledgments

The authors are grateful to Genoanalytica Ltd. (www.genoanalytica.ru, accessed on 18 December 2021, Moscow, Russia) for conducting the technological part of the experiment and the initial statistical analysis. The authors express their deep gratitude to Dr. Nikolin V.P. for participation in the preparation of the experiment, also to Dr. Redina O.E. for the fruitful discussion of the results obtained and Dr. Dyuzhikova N.A. for organizational support.

## Conflicts of Interest

The authors declare no conflicts of interest.

## Abbreviations

C, Cont: control, intact mice
D, Los: depressed mice, losers
DT, Los+Tum: depressed mice after injection of Lewis lung adenocarcinoma cells
DT, Los+Tum*: depressed mice after injection of Lewis lung adenocarcinoma cells with maximal changes of gene expression
DEGs: differentially expressed genes

## Notes

### Competing Interest Statement

The authors have declared no competing interest.

### Funding

- State Funding of the Pavlov Institute of Physiology, RAS: 1021062411629-7-3.1.4

